# Does Tobacco Abstinence Decrease Reward Sensitivity? A Human Laboratory Test

**DOI:** 10.1101/128744

**Authors:** John R Hughes, Alan J Budney, Sharon R Muellers, Dustin C Lee, Peter W. Callas, Stacey C Sigmon, James R Fingar, Jeff Priest

## Abstract

**Introduction:** Animal studies report abstinence from nicotine makes rewards less rewarding; however, the results of human tests of the effects of cessation on reward sensitivity are mixed. The current study tested reward sensitivity in abstinent smokers using more rigorous methods than most prior studies.

**Methods:** A human laboratory study compared outcomes for 1 week prior to quitting to those during 4 weeks post-quit. The study used smokers trying to quit, objective and subjective measures, multiple measures during smoking and abstinence, and monetary rewards to increase the prevalence of abstinence. Current daily smokers (n = 211) who were trying to quit completed an operant measure of reward sensitivity and a survey of pleasure from various rewards as well as self-reports of anhedonia, delay discounting, positive affect and tobacco withdrawal twice each week. A comparison group of long-term former smokers (n = 67) also completed the tasks weekly for 4 weeks. Primary analyses were based on the 61 current smokers who abstained for all 4 weeks.

**Results:** Stopping smoking decreased self-reported pleasure from rewards but did not decrease reward sensitivity on the operant task. Abstinence also decreased self-reported reward frequency and increased the two anhedonia measures. However, the changes with abstinence were small for all outcomes (6-14%) and most lasted less than a week.

**Conclusion:** Abstinence from tobacco decreased most self-report measures of reward sensitivity; however, it did not change the objective measure. The self-report effects were small.

**Implications:**

- Animal research suggests that nicotine withdrawal decreases reward sensitivity. Replication tests of this in humans have produced inconsistent results.
- We report what we believe is a more rigorous test
- We found smoking abstinence slightly decreases self-reports of reward sensitivity but does not do so for behavioral measures of reward sensitivity

## INTRODUCTION

When animals are administered nicotine chronically and this is stopped, then during withdrawal, the animals are less willing to work for rewards ^1-5^ as indicated by increased thresholds for intracranial self-stimulation during abstinence. These effects could represent an “offset” or a “withdrawal effect”^6^. In terms of the former, acute doses of nicotine increase the willingness of animals to work for drug and non-drug rewards^7-9^ and this appears to be true in humans ^8 10, 11^; thus, when smokers quit and lose this effect, they should experience decreased reward sensitivity, simply due to the loss of the direct effects of nicotine; i.e., independent of any neural compensation.. This “offset effect” should produce a gradual unilateral change over time. On the other hand, several studies have labeled the decreased sensitivity as a “withdrawal effect. If this is correct, then deprivation should produce a transient change resulting in an inverted U shaped time course. Whether animal studies indicate an offset or withdrawal effect is unclear, in part, because few have measured reward sensitivity on multiple occasions over time

Several human studies have directly, or indirectly measured reward sensitivity with nicotine abstinence. These studies usually either measure a) a behavioral outcome in which participants either work less hard to obtain presumably rewarding stimuli (e.g. money or preferred music) or, in choice situations, allocate less responding to higher magnitude or more probable rewards or b) a self-report measure of the enjoyment or frequency of rewards or an anhedonia scale. We located 19 such trials ^5, 10-26^ and two additional studies ^13, 16^ that included only self-report outcomes. Most of the above studies used experimental within-participant designs (52%), dependent smokers (67%), overnight abstinence (67%), and smokers not trying to quit (85%). Overall, eight studies had positive results and 13 had null results or results that varied across dependent variables. These mixed results could be due to one or more of the following methodological decisions: a) use of smokers who are not trying to quit for good ^27^, b) only overnight abstinence, c) small sample sizes, d) only one test during abstinence, e) confounding of measures from behavioral tasks by learning/practice effects, f) use of insensitive measures (e.g. only 1-3 questions), and g) outcomes measured well-after the usual time course for withdrawal effects ^28^. The current study attempted to provide a more valid test of abstinence-induced reward sensitivity in humans by minimizing these problems. In addition, our study was designed to determine whether any effects of abstinence appeared to be due to the simple offset of drug effects or due to drug withdrawal. Our major hypotheses were that abstinence would a) decrease reward sensitivity on a behavioral task, and b) decrease ratings of enjoyment from rewards.

## METHODS

### Study Design

We recruited 211 smokers who were trying to quit for good to a study in which they attended two sessions/week for 5 weeks. In the first week, they smoked their usual number of cigs/day. They then quit smoking and were to remain abstinent for 4 weeks. To obtain an adequate number of continuously abstinent smokers to decrease selection bias, we used a schedule of escalating monetary incentives to encourage abstinence. The two primary outcomes were performance on a behavioral task that measures reward sensitivity, and a scale that measures enjoyment from various events/activities. We also recruited a comparison group of 67 long-time former smokers to a) determine whether measures in abstinent smokers have returned to a level similar to long term abstinence and, b) whether repeated testing influences our outcomes. The study occurred at the University of Vermont and Dartmouth College and was approved by the ethics committees of both sites. The study was registered at www.clinicaltrials.gov (NCT01824511).

### Participants

Potential participants were recruited by flyers (22%), Craigslist (www.craigslist.com) (19%), newspaper ads (18%), word-of-mouth (14%), radio ads (11%), and other sources. Generic inclusion criteria for both current and former smokers were a) ≥18 years old, b) able to read and understand English, c) no current (last year) mood or non-nicotine alcohol/drug-related psychiatric disorder, nor any neurological condition that could influence reward sensitivity; e.g. Parkinsonism ^29, 30^, d) no current use of psychoactive medications; e.g. antidepressants or anxiolytics, e) used marijuana ≤ 2 times in the last month, f) agree to no use of non-cigarette tobacco, non-tobacco nicotine, marijuana, illegal drug, electronic cigarettes, or smoking cessation products during the study, and g) not currently pregnant.

The inclusion criteria for the current smoker condition were a) currently smoke <u>≥</u>10 cigarettes daily for <u>≥</u> 1 year, b) want to quit smoking for good via abrupt cessation without treatment, c) willing to quit 7-14 days from study entry and not reduce before quitting, e) have carbon monoxide (CO) level ≥ 8 ppm at consent. We included only those wanting to quit to increase generalizability and sensitivity^27^ and because such smokers have more withdrawal when they quit ^31^ The most common reasons for exclusion were use of a psychoactive medication, use of cannabis and too few cigarettes/day (Figure 1). Just over half of those eligible consented and entered the study (n = 211). Current smokers were generally similar to US current smokers who had recently tried to quit or were planning on quitting in gender, race, age, cigarettes/day and Fagerstrom Test for Cigarette Dependence ^32^ but were more educated (Table 1).

**Table 1.** Comparison of study sample with a population based sample

|  | Current Study |  |  |  |
| --- | --- | --- | --- | --- |
|  | Current Smokers<br>(*n* = 211) | Former Smokers<br>(*n* = 67) | US Current Smokers with Recent Quit Attempt<sup>a</sup> | US Former Smokers<sup>a</sup> |
| % Women | 45% | 46% | 46% | 46% |
| % White/Non-Latino | 86% | 85% | 74% | 74% |
| Age (*M* ± *SD*) | 40 ± 15 | 47±16 | 40 | 40 |
| % Some college or more | 67% | 93% | 46% | 57% |
| % Employed | 62% | 69% | - | - |
| Cigarettes per day (*M* ± *SD*) | 19 ± 8 | - | 16 | - |
| FTCD Total (*M* ± *SD*) | 5.0 ± 2.1 | - | 4.4 | - |

<sup>a</sup>Based on 2007 National Health Interview Survey <sup>74</sup>, National Epidemiological Survey on Alcohol and Related Conditions <sup>75</sup>, and Fagerstrom & Furburg <sup>76</sup> <sup>32</sup> M= mean, SD= standard deviation, FTCD= Fagerstrom Test for Cigarette Dependence

**Figure 1.**
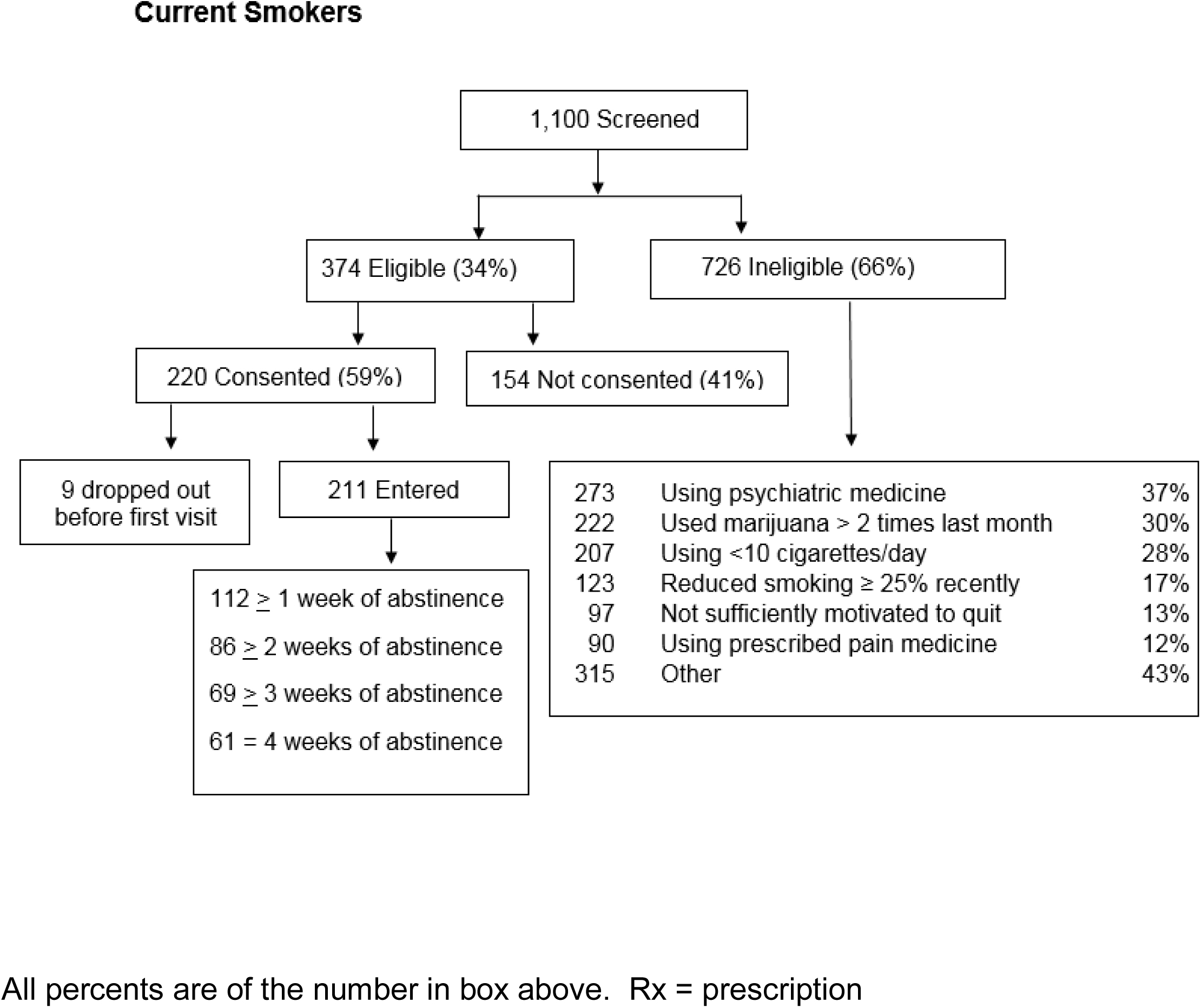
Participant flowchart.

The inclusion criteria for the former smoker condition were a) smoked ≥10 cigs/day for ≥1 years in past, b) used ≤ 5 cigarettes in last year, and c) have not used any non-cigarette tobacco or nicotine products in the last month. Our sample of former smokers was similar to the US average former smoker except for a higher educational level. Former smokers were similar to current smokers except they were older (Wilcoxon rank sum test Z = 3.4, p < 0.001) and more educated (x^2^= 49.9, p < 0.001).

### Sample Size

Our original aim was to recruit until we obtained 70 smokers who abstained for 4 weeks. This sample size would provide > 90% power to detect a change of 20% in scores on most of our dependent variables after abstinence, if we assumed a within-participant correlation of 0.8, which is similar to that found in our prior withdrawal studies ^31^.

### Procedures

We used an escalating payment schedule to increase abstinence rates. Participants attended two laboratory visits/week to provide breath samples for CO and urine samples for cotinine to verify self-reported abstinence. A CO of < 8 ppm (Smokerlyzer, Bedfont) at both visits in the first week of instructed abstinence was required to assume initial abstinence. Although twice daily CO would be necessary to truly verify smoking, a recent study found daily or almost daily CO validation to be an adequate substitute ^33^. For the second through fourth weeks of abstinence, we added a criteria that the urine cotinine test strip (Onescreen cotinine test, American Screening) have a value = 0 indicating cotinine < 10 ng/ml. These CO and cotinine cutoffs detect recent smoking/abstinence with high sensitivity and specificity ^34^. We also required a negative urine cannabis dipstick result (Discover THC dipstick, American Screening) because cannabis use might mimic decreased reward sensitivity^35^. The monetary reward schedule was similar to that effective in our prior studies ^36, 37^. The abstinent-contingent payments began at $16/visit and increased at each subsequent visit to a maximum of $30/visit. In addition, participants could receive $50 - $100 bonuses payments for continuous abstinence. Participants abstinent for all 4 weeks received $534. Research staff provided supportive counseling at each visit (about 5 minutes) consistent with the USPHS guidelines ^38^.

Former smokers attended lab visits once a week for 4 weeks. Tobacco and cannabis abstinence were verified as with current smokers; however, we did not provide extra payment for abstinence. Both current smokers and former smokers also received payments for attending visits and completing measures and these payments were not contingent on not smoking.

### Measures

One primary measure was the Effort Expenditure for Rewards Task (EEfRT) that examines responding as a function of response cost, reward magnitude, and probability of reward ^39^. The task presents participants with repeated choice tests. At each test, the program presents a choice between a more difficult task in which success is rewarded with more money, or a less difficult task in which success is rewarded with less money. Participants had 3 seconds to choose which task to undertake. The harder task required 100 button presses with the non-dominant little finger in 21 seconds. The easy task required 30 presses within 7 seconds. The entire session lasted 20 minutes. The payment for each test was randomly assigned to vary from $0.25 - $1.05 and the probability of payment was either 12%, 50% or 88%. Participants were informed on the payment and probability for each task prior to making a choice. The original EEfRT also includes a second varying probability for whether there is any payment for the test session. To keep the task easier to understand, we deleted this last probability. Reward responsivity was measured by the proportion of higher reward tasks chosen across all probabilities and then separately for the high, medium and low probability choice tests. Decreased choice of the higher reward test would indicate decreased reward sensitivity. Of the 87,787 trials, we excluded 2030 (2.3%) because the participant did not make a choice of hard vs easy, or exclusively chose hard or easy throughout the session. Performance on the EEfRT has been shown to be correlated with self-report measures of anhedonia and is sensitive to the effects of stimulants to increase reward seeking ^39-41^

The other primary measure was the Rewarding Events Inventory, a self-report measure of 54 common rewards that we developed to be a more comprehensive and up-to-date measure than existing scales. The measure has excellent internal validity and test-retest reliability ^42^. The Inventory asked participants to rate the events separately on enjoyment, with response options of “not enjoy it, enjoy it a little, enjoy it some, enjoy it a lot, extremely enjoy it, and frequency, with response options of every day, most days, few days, one day or no days in the last week.”

To verify that participants were having withdrawal symptoms during abstinence we asked participants to rate the nine DSM-5 (www.dsm5.org) withdrawal items from 0= not at all to 4=severe (nb, this does not include craving), using the Minnesota Nicotine Withdrawal Scale-Revised (www.uvm.edu/∼hbpl) ^28, 43^. The MNWS has good psychometrics ^28, 43^. We also included measures of constructs related to reward sensitivity; i.e., anhedonia/apathy, delay discounting, and positive affect to provide convergent validity tests. One measure was the 18-item Apathy Evaluation Scale (AES) which asks about decreased pleasure or interest in rewards (e.g. “I am interested in having new experiences”) ^44^ with ratings from 1 = not at all to 4 = very true. The other was the Temporal Experience of Pleasure Scale (TEPS) that includes a 10 item interest in reward scale and an 8 item pleasure from reward scale with response options on a 6 point Likert scale ^45^. The AES and TEPS have been tested in psychiatric, drug abuse and neurological patients, and have good reliability and adequate validity ^46-48^. Positive affect was measured using the Positive and Negative Affect Scale (PANAS) ^49^. This widely used measure has excellent reliability and validity ^50^. We also examined delayed discounting (DD) because decreased reward sensitivity appears to be associated with decreased delay discounting ^51^. For this measure, participants completed a delay discounting task ^52^ in which participants are given a series of hypothetical choice situations of receiving money now vs later with varying monetary values and delay periods. The amount available after the delay was always $1000 and the range of delays was 1 day to 5 years. Across trials, a smaller, sooner reward was adjusted up or down by 50% depending on the subject’s choice (smaller, sooner choices resulted in decreases; larger, delayed choices resulted in increases).

Results with these hypothetical choices are consistent with actual choices ^53^. The major outcome was the k statistic which reflects the relative preference for a small, immediate reward vs. a larger delayed reward and is based on a natural log transformation of k ^52^. A decrease in preference for the more immediate reward would assumted to indicate decreased reward sensitivity. All of the above measures were obtained at every lab visit.

In summary, we expected that abstinence would increase withdrawal scores, decrease choice of hard response on the EEfRT, decrease enjoyment and frequency of rewards on the REI, increase anhedonia scores on the AES and TEPS, decrease positive affect, and decrease delay discounting.

### Data Analysis

Since many smokers do not maintain abstinence during a withdrawal study, the major issue in analysis is whom to include in the analyses^28^. If one uses all participants, this reduces bias and generalizability resulting from examining only a subset of participants, but it requires using withdrawal scores among those who currently smoke. If one uses only those abstinent for the entire study, this avoids using smokers who are smoking but can substantially reduce the sample size and allow selection bias. Another option is to include all smokers and use abstinence status as a time-varying covariate^54^. This option includes all smokers but some of the withdrawal scores are based on short periods of abstinence. For the primary analysis we chose to use the 61 participants who were abstinent for the entire four weeks for four reasons: a) this is the most commonly used option in tobacco withdrawal studies^31^, b) several days of abstinence may be necessary to change reward sensitivity, c) our resultant sample size was still adequate for our within-participant analyses, and d) this allows a test of time pattern (i.e., whether the results appear to be due to simple drug offset or to drug withdrawal) that is not influenced by different participants at different timepoints. We also undertook two sensitivity tests using a) the larger sample (n= 104) of participants abstinent during the first week when abstinence symptoms typically peak, and b) all 211 participants with abstinence status as a time-varying covariate.

Our major analyses were based on within-participant ANOVAs. We used mixed linear modeling to conduct longitudinal analyses of outcomes, including both restricted maximum likelihood of fixed effects with a compound symmetric covariance matrix, and random effects with an unstructured covariance matrix. All analyses were conducted using SAS 9.4 software (SAS Institute, Cary, NC) and statistical significance across all tests was defined as p < .05 (2-tailed). Our pre-specified major test among current smokers was a comparison of the mean EEfRT and REI enjoyment scores during the smoking-as-usual period vs the mean during the entire abstinence period. We also specifically tested for an inverted U shape pattern in abstinence for all outcomes; i.e., whether any outcomes had any initial increase/decrease or during abstinence which abated over time during the abstinence period via an ANOVA confined to the abstinence period. We also used paired t tests to compare peak baseline score and peak score during abstinence using the highest score as peak for those measures expected to increase with abstinence and lowest score for those expected to decrease. We next compared results between former smokers and newly abstinent smokers to see if the recent abstainers had returned to the level of long-term former smokers. To do this, we tested whether the results at the 3^rd^ and 4^th^ visits among abstainers differed from the results from the 3^rd^ and 4^th^ visit results among long-term former smokers, again with an ANOVA.

## RESULTS

### Initial Analyses

About half (n=104, 52%) of participants were abstinent for > 1 week but this decreased to about a fourth (n = 61, 29%) abstinent for all 4 weeks (Figure 1; Appendix, Figure 1). As in most clinical studies, those who were able to abstain longer (i.e., for 4 weeks) were older (Wilcoxon rank sum test Z = 2.0, p = .05), more educated (Fisher’s Exact Test, p = .004) and smoked fewer cigs/day (Wilcoxon rank sum test Z = 3.0, p =.003) than those who were not able to do so. Among the 61 fully abstinent participants, the two baseline values did not differ from each other for any outcome, indicating they represent stable baseline scores. The mean score for the MNWS score during abstinence was substantially greater than the mean score during smoking (+43%) indicating these 61 participants were in withdrawal during the abstinence period (F=64.8, p< 0.001). For most measures, the scores for the four tests among long-abstinent smokers found little change with repeated testing (< 2.5% change from one time point to the next); however, preference for the harder task on the EEFRT increased overtime (6.7% increase between tests). There were no significant differences between the 3^rd^ and 4^th^ visits for long-term former smokers.

### Main Analyses

: Contrary to our hypothesis, the mean proportion of choices that were for the higher-reward task on the EEfRT task during the abstinence period was greater, not smaller, than the mean proportion during the smoking period (F=40.4, p < 0.001; see Table 2, Figures 2 and 3). When we looked at results at each of the three probability of reward settings on the EEfRT, one showed no change and the other two showed an increase, not a decrease; at probability =0.12, F=0.7, p =0.39; at probability=0.50, F= 52.2, and p<0.001; at probability=0.88, F=82.3, p<0.001. In contrast, consistent with our major hypothesis, abstinence decreased the rated enjoyment from rewards on the REI, F=133.1, p<0.001. Also, consistent with our hypotheses, abstinence decreased the frequency of rewarding events, (F=58.4, p < 0.001) and the delay discounting outcome, (F=22.5, p<0.0010) and increased scores on the AES, F=11.4, p=0.001, and TEPS scales, (F=5.5, p=0.02), However, the magnitudes of change for both our primary and secondary outcomes were small (6-14%, see Table 2 and Figures 2 and 3). Using the mean score across all abstinence measures as the dependent variable could have obscured a change that occurred on only one or two days during abstinence. To test this, we reran the analyses comparing the peak value during abstinence vs the peak value during smoking. The results were very similar (Appendix Table 1).

### Time Course

True withdrawal symptoms exhibit an inverted U time course; to test this we examined whether, among the five outcomes that showed the hypothesized initial change with abstinence (REI enjoyment, REI frequency, DD, TEPS and AES), whether these changes scores then decreased over time (i.e., we compared scores in week 1 vs the average across weeks 2-4). This was true for the REI enjoyment (t = 2.1, p = 0.04), REI frequency (t = -3.2, p = 0.002) and DD (t = -2.4, p =.02) outcomes but not for the AES or TEPS scores. Figures 2 and 3 illustrate the magnitude of these results

**Table 2.** Baseline vs. Abstinent Scores in Participants Abstinent for Four Weeks (n=61) and Among Positive Results, Whether Time Course Was Consistent With That of a Withdrawal Symptom

| Mean Baseline vs. Mean Abstinent |  |  |  |  |  |
| --- | --- | --- | --- | --- | --- |
| Variable (range) | Baseline | Abstinent | Absolute Change | Percent Change | Withdrawal Time Pattern<sup>b</sup> |
| EEfRT (0-1) | 0.51 | 0.58***<sup>a</sup> | +0.07 | +14% | No |
| REI Enjoy (1-5) | 3.7 | 3.5*** | -0.2 | -5% | Yes |
| REI Frequency (1-5) | 2.6 | 2.4*** | -0.2 | -8% | Yes |
| AES (18-72) | 29.2 | 30.9*** | +1.7 | +6% | No |
| TEPS (18-108) | 41.7 | 43.5* | +1.8 | +4% |  |
| PANAS PA (10-50) | 33.8 | 33.3 | -0.5 | -2% |  |
| Delay discounting (ln k) | -6.4 | -6.8*** | +0.4 | +6% | Yes |

<sup>a</sup>Note opposite to hypothesized change
\* p < .05
\*\* p < .01
\*\*\* p < .001
AES = Apathy Evaluation Scale, EEfRT = Effort Expenditure for Rewards Task, K = test statistic for degree of discounting, ln = natural log, NA = Negative Affect, PA = Positive Affect, PANAS = Positive and Negative Affect Scale, REI = Rewarding Events Inventory, TEPS = Temporal Experience of Pleasure Scale
<sup>b</sup>Withdrawal pattern defined as first post-cessation value greater than remaining post-cessation values

**Figure 2.**
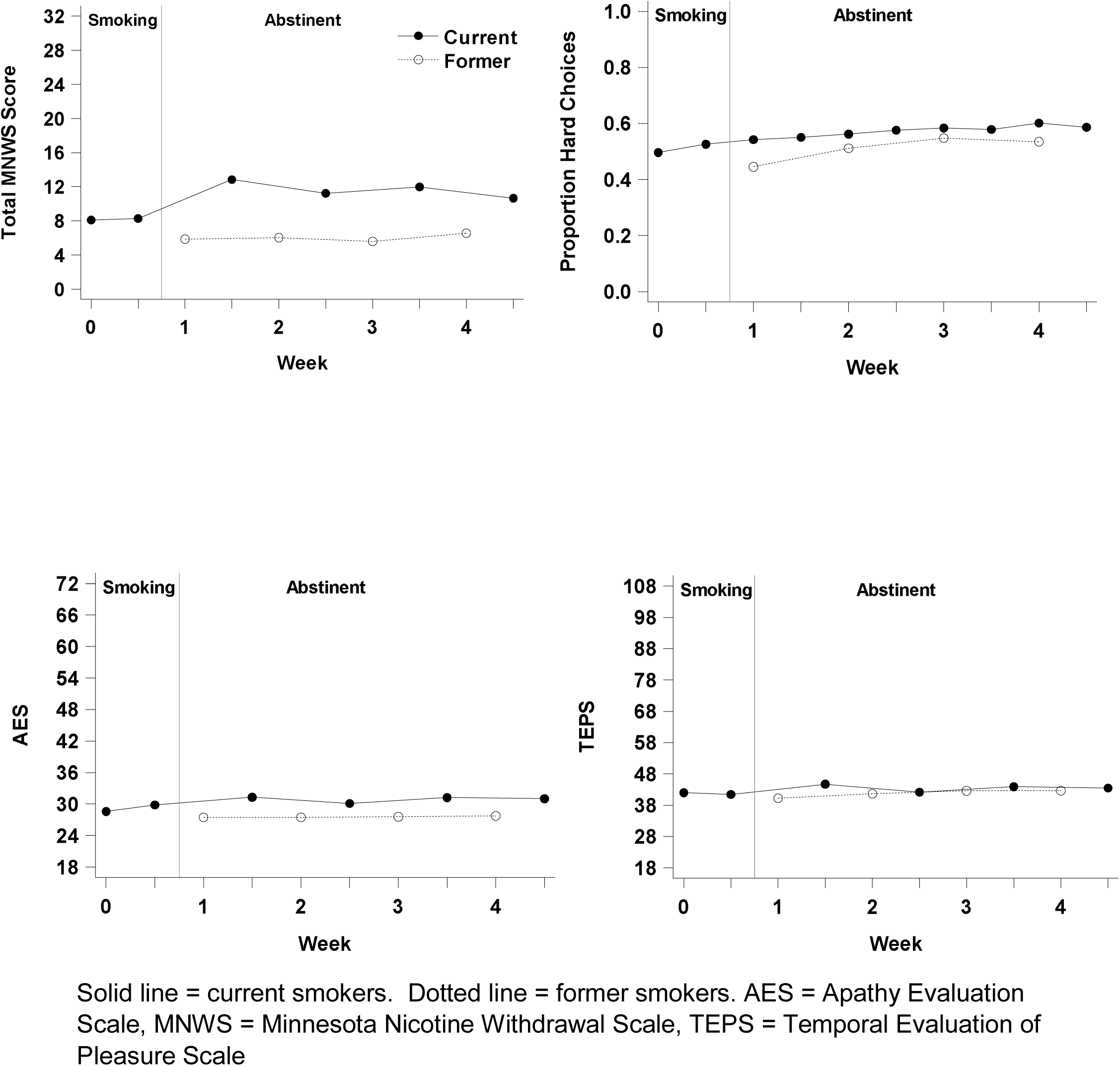
Time course for selected outcomes.

**Figure 3.**
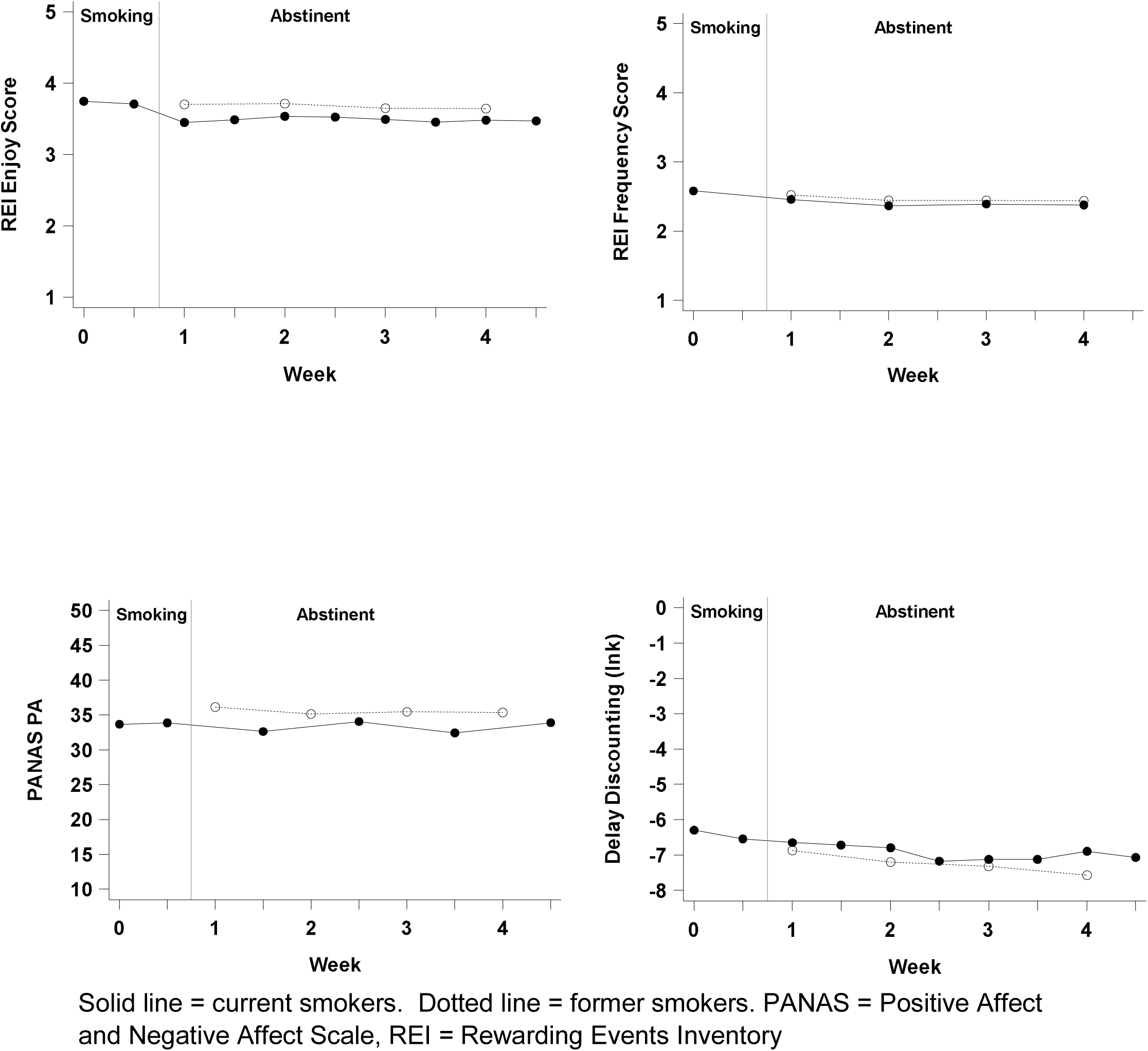
Time course for selected outcomes.

### Former Smokers

After adjusting for subject characteristics that differed between former and current smokers, recently abstinent smokers had higher MNWS scores, (F = 30.1, p < 0.001), and AES scores, (F = 4.4, p = .04); but lower positive affect scores, (F = 8.5, p < 0.004), than former smokers. Abstinent smokers also had lower REI enjoyment scores (Wilcoxon rank sum test Z = -2.1, p = .03), although we could not meet ANOVA assumptions for including baseline differences in this particular test. Figures 2 and 3 illustrate the magnitude of differences.

### Sensitivity Analyses

We reran the major analysis; i.e., a comparison of mean smoking vs mean abstinence score for the EEfRT and REI enjoyment but only examined results from the first week of abstinence and included all participants abstinent during the first week (n = 104). We also reran analyses using all participants (n = 211) and using abstinent state as a time-varying covariate. In both analyses, the results were very similar to that for the 61 long-abstinent smokers. The results of these analyses are in Appendix Tables 2 and 3..

## DISCUSSION

Cigarette abstinence decreased self-reports of pleasure from and frequency of rewards on a reward inventory, and increased scores on the two anhedonia scales. That all four of these changed in the hypothesized direction suggests convergent validity of results. On the other hand, the magnitude of change in these outcomes was only 4-8%. Among the 21 prior studies of the effect of abstinence on reward sensitivity, 6 examined change in self-reports, among these, 3 found anhedonia increased but only one reported the magnitude of change (+19% estimated from graph)^16^ and this was probably the study with the highest internal and external validity.

Abstinence did not decrease reward sensitivity in the behavioral test of reward sensitivity - the EEfRT. Among the 15 prior studies that examined a behavioral task of reward sensitivity, 7 (47%) reported abstinence decreased sensitivity. Among the six studies that reported magnitude of effects, the median decrease was only 2.5%. However, the study that was the most rigorous test and used a validated behavioral task found abstinence completely eliminated preference for the higher magnitude reward ^5^.

One possible reason we failed to find an effect on EEfRT on the behavioral task was that scores on the EEfRT appeared to increase over time, suggesting a learning effect that may have obscured any decline due to abstinence. Prior studies have not measured the EEfRT repeatedly over time; thus, whether learning effects are common with EEfRT requires further testing. Another possible reason for different outcomes in self-report vs behavioral task outcomes is that they are measuring two different aspects of reward sensitivity ^55^; i.e., the former is measuring hedonic response and the latter motivation to pursue rewards.

In terms of other secondary outcomes, if abstinence decreases reward sensitivity, then this should decrease delay discounting ^56^ which is what we found, albeit the magnitude of this effect was small. Across the ten prior studies of the effect of initial abstinence on delay discounting, two found abstinence decreased DD ^52, 57^, four found it produced no change ^33, 58-60^ and four found abstinence increased DD ^61-64^. Our reading of these studies does not suggest a clear reason for these heterogeneous results. It may be because the DD task and how its results are calculated differs across studies. Also, studies that found abstinence increased DD, interpreted this as indicating abstinence increases impulsivity, which is consistent with other studies ^65^. Thus, it may be that abstinence has two opposing effects on DD tasks; i.e., it decreases reward sensitivity but also increases impulsivity.

One possible limitation of our results is that our main analysis used only the 30% of our participants who were abstinent for the desired 4 week period; however, tests using a larger sample of those initially abstinent and using all participants obtained similar results. Also, we did not conduct a randomized trial of abstinent vs non-abstinent conditions. Instead we used a pre-post design in which participants served as their own control. Although such non-randomized designs can have methodological problems, in fact, most studies of tobacco withdrawal have used pre vs post designs and have provided very replicable data. We did not include several measures of reward sensitivity such as neuroimaging ^66^ or response to hedonic stimuli ^19, 67^ that tap other aspects of reward sensitivity; e.g. reward anticipation or reward learning. Our participants were more educated and more nicotine dependent than the average US smoker and had no use of psychoactive drugs or current psychiatric disorder; this may decrease the external validity of our study. The influence of abstinence may differ by psychiatric status^5^; however, we did not collect information on past or current psychiatric status. Our sample size for analysis might be thought of as small; however, our data analysis was based on a within-participant comparison with multiple pre and post-cessation values.

To our knowledge, this was the first use of the EEfRT to examine the effects of smoking abstinence. Several studies have found EEfRT can detect reward sensitivity changes with other drugs ^39, 41, 68-71^. Our first post-cessation EEfRT measurement did not occur until 3-4 days after cessation. Some prior studies suggest the effect of abstinence on reward sensitivity occurs immediately after cessation ^5^; thus we could have missed a short-lived effect. Although one could believe such a “fleeting” effect would not be clinically important, this may not be the case because over half of all relapses occur in the first 3 days^72^. Finally, our contingency program to increase abstinence relied on testing at only 2 visits/week and thus, some smoking could have occurred between the visits and this decreased the sensitivity of our test. On the other hand, the scores on the MNWS increased substantially indicating participants were in withdrawal during the last 4 weeks of the study.

Compared to the prior studies of the effect of tobacco abstinence on reward sensitivity, the current study had the following methodological assets: a) use of smokers who are trying to quit for good, b) longer period of abstinence that included the typical time periods for tobacco withdrawal, c), a sufficient time period to detect if effects due to withdrawal or simple offset of direct effects, d) multiple measures during smoking and abstinence periods, e) use of both objective and subjective measures, f) multiple measures to allow test of convergent validity, g) abstinence verification on multiple occasions, and h) experimental induction of abstinence. ^28^

In summary, our self-report results suggest abstinence induces a small decrease in reward sensitivity, but our behavioral task did not confirm this. Animal studies clearly predict that smoking cessation should decrease reward sensitivity (see above). We believe the small decrease in reward sensitivity we observed in a subset of the measures is a weak confirmation of the applicability of animal data to smoking cessation in humans. Given the heterogeneity of prior results on whether smoking abstinence decreases reward sensitivity and the limitations of our own methods, further studies of abstinence-induced decreased reward sensitivity using other behavioral tasks such as the Progressive Ratio task ^10^ or the Signal Detection Task ^73^ that may be more sensitive or reliable, concomitant with self-report measures, are needed to further clarify the importance of reward sensitivity to smoking cessation.

## Funding

This study was supported by a grant from the US National Institutes of Health (1 R01 DA-031687).

## Acknowledgements

We thank Jean Francois Etter for help in designing the study and in conducting studies developing the Rewarding Events Inventory. We thank Michael Treadway for sharing his EEfRT task with us. We also thanks Bonita Basnyat, Erin Kretzer, Doris Gasangwa, Grey Norton, Stanley See, Hao Yani for assistance in conducting the study and Jessie McNabb for help in preparing the manuscript..

## Declaration of Interests

JRH has received consulting fees from companies that develop or market products for smoking cessation or harm reduction and from non-profit companies that engage in tobacco control. Other authors have nothing to disclose.

